# Your Brain Doesn’t Look a Day Past 70! Cross-Sectional Associations with Brain-Predicted Age in the Cognitively-Intact Oldest-Old

**DOI:** 10.1101/2025.05.26.655855

**Authors:** Mark K. Britton, Hannah Hoogerwoerd, Joshua Juhasz, Keyanni Joy Johnson, Paul D. Stewart, Stacy S. Merritt, Cortney J. Jessup, Clinton B. Wright, David A. Raichlen, G. Alex Hishaw, Victor A. Del Bene, Virginia G. Wadley, Theodore P. Trouard, Noam Alperin, Bonnie E. Levin, Tatjana Rundek, Kristina M. Visscher, Gene E. Alexander, Ronald A. Cohen, Eric C. Porges, Joseph M. Gullett

**Author notes:** Corresponding Author: Mark K. Britton, Department of Epidemiology, University of Florida, Gainesville, FL.

## Abstract

The cognitively-intact oldest-old (85+) may be the most-resilient members of their birth cohort; due to survivorship effects (e.g., depletion of susceptibles), risk factors associated with brain aging biomarkers in younger samples may not generalize to the oldest-old. We evaluated associations between established aging-related risk factors and brain-predicted age difference (brainPAD) in a cross-sectional cognitively-intact oldest-old sample. Additionally, we evaluated brainPAD-cognition associations to characterize brain maintenance vs. cognitive reserve in our sample. Oldest-old adults (N = 206; 85-99 years; MoCA > 22 or neurologist evaluation) underwent T1-weighted MRI; brainPAD was generated with brainageR, such that more-positive brainPAD reflected relatively advanced brain aging. Sex, educational attainment, alcohol and smoking history, exercise history, BMI, cardiovascular and metabolic disease history, and anticholinergic medication burden were self-reported. Global cognitive z-score and coefficient of variation were derived from the NACC UDS 3.0 cognitive battery; crystallized-fluid discrepancy was derived from the NIH Toolbox Cognitive Battery. Mean brainPAD was -7.99 (SD: 5.37; range: -24.50, 6.03). Women showed more-delayed brain aging than men (B = -2.35, 95% CI = - 4.28, -0.41, p = 0.018). No other exposures were associated with brainPAD. BrainPAD was not associated with any cognitive variable. These findings suggest that cognitively-intact oldest-old adults may be atypically-resistant to risk factors associated with aging in younger samples, consistent with survivorship effects in aging. Furthermore, brainPAD may have limited explanatory value for cognitive performance in cognitively-intact oldest-old adults, potentially due to high cognitive reserve. Overall, our findings highlight the impact of survivorship effects on brain aging research.

**Highlights:**

- Brain-predicted age difference was assessed in cognitively-intact oldest-old (≥85)
- Mean brain-predicted age corresponded to an 8-year delay in brain aging
- Brain age in oldest-old was not associated with self-reported health history
- Brain age was not associated with cognitive performance

## 1. INTRODUCTION

Oldest-old adults (≥85 years) are, by definition, survivors among their birth cohort. An estimated 23% of men and 38% of women born in the United States in 1930 reached the age of 85 (Bell and Miller, 2005). An even smaller subset of individuals maintain largely-intact cognitive function into oldest-old age: the 2022 National Health Interview Survey reported that 13.1% of noninstitutionalized US adults 85 years and older had ever been diagnosed with dementia (Kramarow, 2024), while estimates of Mild Cognitive Impairment (MCI) burden in noninstitutionalized adults 80 years and older range from 16-27% (Bai et al., 2022). However, a substantial minority of oldest-old adults do maintain intact cognitive performance, whether due to physiological resistance to aging (brain maintenance) or strong compensatory abilities (i.e., cognitive reserve) (Kawas et al., 2021; Stern, 2009). Consequently, a growing body of research has sought to identify protective factors associated with brain function in cognitively-intact oldest-old adults, with the goal of identifying potential interventions to prolong cognitive function in aging (Andersen, 2020; Silverman and Schmeidler, 2018).

Brain-predicted age is an emerging biomarker of overall physiological brain aging (Cole and Franke, 2017; Franke and Gaser, 2019). Brain-predicted age refers to a chronological age value predicted from a structural brain image (e.g., a T1-weighted MRI) using supervised machine learning regression. Brain-predicted age difference (brainPAD), or the difference between chronological age and brain-predicted age, has been interpreted as a biomarker of advanced vs. delayed brain aging in middle-aged and young-old adults (Cole and Franke, 2017). brainPAD is associated with midlife exposures linked to physiological aging, such as physical activity and fitness (Bittner et al., 2021; Dunås et al., 2021; Steffener et al., 2016), cardiovascular (Cherbuin et al., 2021; De Lange et al., 2020; Wagen et al., 2022) and metabolic health (Beck et al., 2022; Franke et al., 2014, 2013; Kolbeinsson et al., 2020; Wing et al., 2022), smoking (Bittner et al., 2021; Linli et al., 2022), and heavy alcohol use (Guggenmos et al., 2017). Furthermore, more-positive brainPAD (i.e., advanced brain age) has been linked to worse cognition in older adults (Cole et al., 2018; Cumplido-Mayoral et al., 2024; Jawinski et al., 2022; Park et al., 2025; Wrigglesworth et al., 2022b) and outperforms total brain volume and MMSE as a predictor of dementia hazard in at-risk older adults (Biondo et al., 2022; Gaser et al., 2013). In sum, brainPAD is a potentially informative biomarker of physiological brain aging and its relationship to cognition in oldest-old adults.

Despite brainPAD’s potential utility in this population, relatively few studies of brainPAD have included oldest-old adults (Boyle et al., 2021; Cumplido-Mayoral et al., 2023; Park et al., 2025; Wrigglesworth et al., 2022b) and, to our knowledge, none have reported data including only oldest-old adults. However, associations reported in young-old adults may not generalize to the oldest-old. Although structural brain aging in oldest-old adults is generally understudied (Merenstein and Bennett, 2022), age effects on brain macrostructure have been suggested to decrease nonlinearly in oldest-old age: for instance, in cross-sectional data, oldest-old adults show weaker age effects on white matter volume (Yang et al., 2016) and sulcal widening (Tang et al., 2021) relative to young-old adults. We have previously reported similar attenuation of age effects on neurotransmitter levels in the oldest-old (Britton et al., 2025), consistent with relative brain preservation in this population.

The apparent attenuation of age effects on brain structure and function in cross-sectional samples of the oldest-old may be partly explained by survivorship effects. Individuals who have survived to oldest-old age are likely atypically healthy or resilient (Borras et al., 2020; Cabeza et al., 2018; Hernán et al., 2008); this is especially true for the cognitively-intact oldest-old. Cognitively-intact oldest-old individuals may be resistant to age-related brain changes (brain maintenance) or more able to compensate for those changes (cognitive reserve), potentially buffering the effects of aging or lifestyle exposures (Arenaza-Urquijo and Vemuri, 2018; Cabeza et al., 2018; Merenstein and Bennett, 2022). Conversely, the individuals most susceptible to age-related or lifestyle-related brain changes are likely to have been eliminated from the sampled population due to earlier mortality or earlier onset of cognitive impairment (i.e., depletion of susceptibles) (Hernán et al., 2008). Attrition of less-healthy individuals may therefore attenuate associations between risk factors and brain structure and function with increasing age (Legdeur et al., 2018); for instance, associations between diabetes and dementia (Corrada et al., 2008), as well as APOE4 and dementia (Corrada et al., 2013; Juva et al., 2000), may not generalize from young-old samples to the oldest-old. Consequently, it is unknown whether exposure-brainPAD associations in young-old adults generalize to the oldest-old.

The current secondary analysis aims to fill this literature gap by assessing exposure-brainPAD associations in a cross-sectional sample of cognitively-intact oldest-old adults. Lifetime substance use, lifetime history of cardiovascular and metabolic disorders, lifetime physical activity, Body Mass Index (BMI), and educational history were selected for analysis due to reported associations with brainPAD in young-old samples; anticholinergic medication burden was selected due to reported associations between anticholinergic use and brain aging (Risacher et al., 2016) as well as cognitive decline and dementia studied both cross-sectionally (Pieper et al., 2020) and longitudinally (Ancelin et al., 2006). We hypothesized that greater lifetime substance use, greater lifetime history of cardiovascular and metabolic disorders, lower lifetime physical activity, greater BMI, lower educational attainment, and greater anticholinergic burden would be associated with an older-appearing brain. As a second aim, to differentiate brain maintenance from compensatory ability in our sample, we examined associations between brainPAD and global cognitive performance, cognitive coefficient of variation (a measure of intra-individual variability, which has been linked to cognitive aging and dementia) (Aita et al., 2024a; Dykiert et al., 2012), and fluid-crystallized cognition discrepancy (an index of cognitive decline) (O’Shea et al., 2018). We hypothesized that a younger-appearing brain would be linked to better overall cognition, more consistent cognitive performance, and smaller fluid-crystallized discrepancy, consistent with successful brain maintenance.

## 2. METHODS

### 2.1 Participants

Oldest-old adults (N = 206) were recruited at four study sites as part of a larger parent study of brain and cognitive health in the oldest-old (McKnight Brain Aging Registry). All participants were 85 years of age or older and were cognitively-intact as determined by either scores of 28 or more on the Telephone Interview for Cognitive Status – Modified (TICS-M) and 22 or more on the Montreal Cognitive Assessment (MoCA) or a neurologist’s evaluation. Study exclusion criteria were the following: major physical disability; medical conditions expected to limit life expectancy or interfere with study participation (e.g., unstable cancer or uncontrolled severe Major Depressive Disorder); current DSM-5 substance use disorder (SUD); inability to independently perform instrumental activities of daily living (IADLs) or basic activities of daily living (ADLs); less than sixth-grade reading level; or any condition that would interfere with study procedures (e.g., hearing or vision loss, MRI contraindications). All participants provided written informed consent. Study procedures were approved by Institutional Review Boards at the University of Florida (201300162), University of Miami (20151783), University of Arizona (1601318818), and University of Alabama at Birmingham (X160113004) and were conducted in accordance with the Declaration of Helsinki.

### 2.2 Demographic and Clinical Variables

Participants self-reported age, assigned sex, race/ethnicity, years of education, and whether they had ever been diagnosed with hypertension, high cholesterol, diabetes mellitus, myocardial infarction, or cardiac arrest. Additionally, participants self-reported history of lifetime alcohol abuse (yes/no), lifetime smoking (yes/no), duration of smoking in years, and number of cigarettes smoked per day (<1/1/2-5/6-15/16-25/26-35/≥35); number of cigarettes/day was estimated numerically using the center of each category (0.5/1/3.5/10/20/30/40) and multiplied by the duration of smoking to generate approximate pack-years. Pack-years were categorized (nonsmoker/< median/≥ median) due to the approximately log-normal distribution and high proportion of nonsmokers. BMI was calculated from self-reported height and weight.

Cumulative lifetime physical activity was assessed using an adapted version of the Lifetime Physical Activity questionnaire (Friedenreich et al., 1998). Hours per week (0/0.5/1/1.5/2/3/4-6/7-10/≥11) and months per year (0/1-3/4-6/7-9/10-12) of moderate and strenuous activity were reported for 9 time periods (high school/18 years/25 years/35 years/45 years/55 years/65 years/74 years/the past 3 years). The midpoint of each hour and month category was taken. Hours per week were multiplied by 4 to represent hours per month, then by approximate months per year. Moderate activity was multiplied by 4.5 and strenuous activity by 8.5 to represent METs (Ainsworth et al., 2011; Maessen et al., 2016), which were summed to reflect cumulative lifetime activity; cumulative lifetime activity was then categorized by quartile, with Q1 representing the least lifetime activity and Q4 the greatest activity.

Participants self-reported all current medications. Current anticholinergic burden was assessed with the 2012 updated version of the Anticholinergic Cognitive Burden Scale (ACB) (Boustani et al., 2008; Campbell et al., 2013). Due to the right-skewness of the distribution, ACB score was binarized for analysis (no anticholinergic medication/anticholinergic medication).

### 2.3 Neuropsychological Testing

Participants completed the Uniform Data Set (UDS 3.0) neurocognitive testing battery (Weintraub et al., 2018). z-scores were generated for each measure using the regression-based UDS 3.0 norms corrected for sex, age, and years of education (Weintraub et al., 2018). The mean z-score across all measures for each participant was used as an index of global cognitive performance. A full list of measures is reported in Table 1.

**Table 1.** Measures taken from the UDS 3.0 neuropsychological battery.

| **Global Cognitive z-Score** | **Coefficient of Variation** | **Domain** |
| --- | --- | --- |
| Craft Story 21 immediate recall, verbatim | Craft Story 21 immediate recall, verbatim | Memory |
| Craft Story 21 immediate recall, paraphrase | - | Memory |
| Benson complex figure copy, total score | Benson complex figure copy, total score | Visuospatial |
| Number span test forward, total correct | Number span test forward, total correct | Attention |
| Number span test forward, longest span | - | Attention |
| Number span test backward, total correct | Number span test backward, total correct | Attention |
| Number span test backward, longest span | - | Attention |
| Animals list generation, total in 60s | Animals list generation, total in 60s | Language (category fluency) |
| Vegetables list generation, total in 60s | Vegetables list generation, total in 60s | Language (category fluency) |
| Trail-Making Test, Part A | Trail-Making Test, Part A | Processing speed |
| Trail-Making Test, Part B | Trail-Making Test, Part B | Executive function |
| Craft Story 21 delayed recall, verbatim | Craft Story 21 delayed recall, verbatim | Memory |
| Craft Story 21 delayed recall, paraphrase | - | Memory |
| Benson complex figure recall, total score | Benson complex figure recall, total score | Visuospatial/Memory |
| MINT total score | MINT total score | Language (naming) |
| Phonemic test, total F-words and L-words | Phonemic test, total F-words and L-words | Language (verbal fluency) |

**Table 2.**
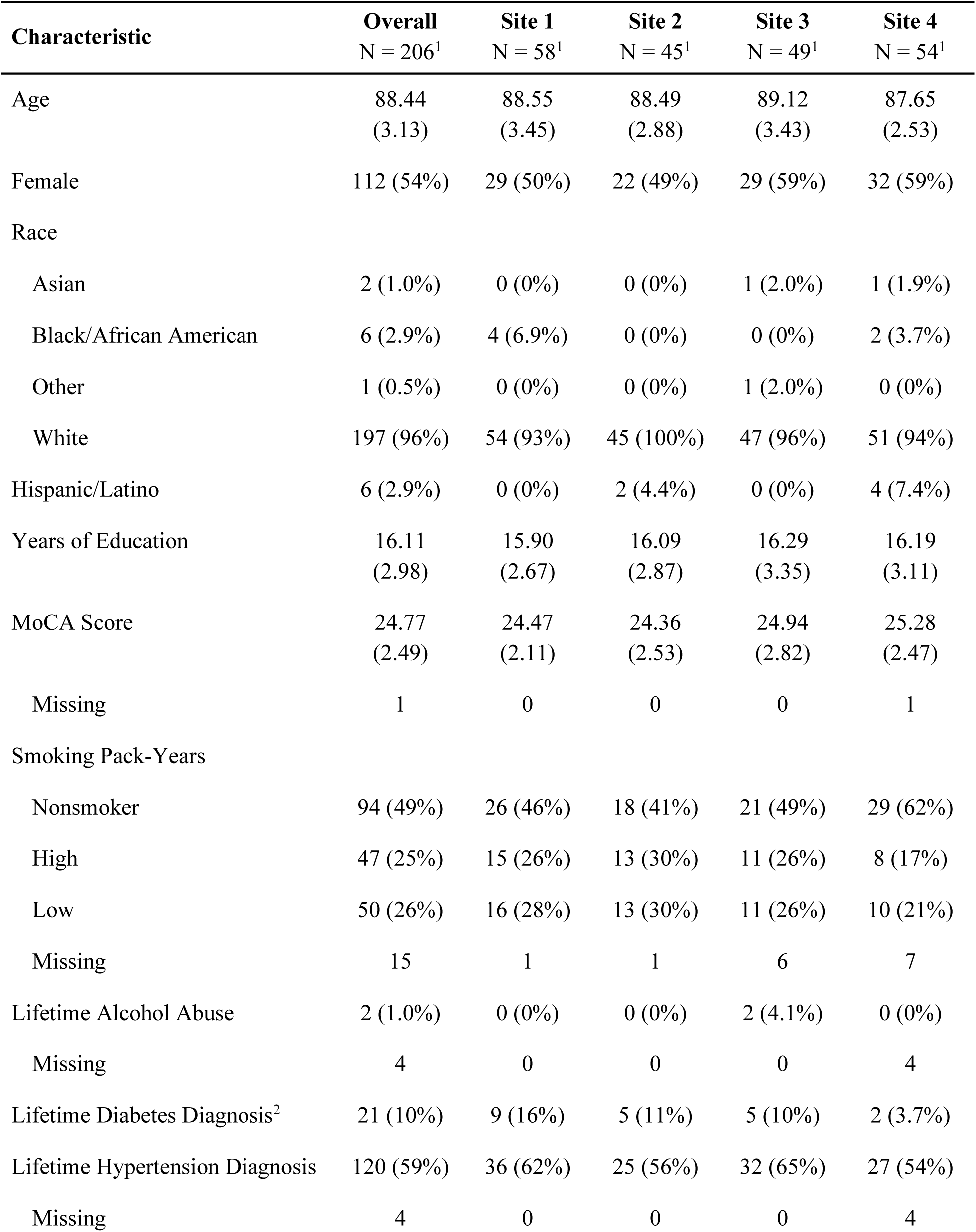
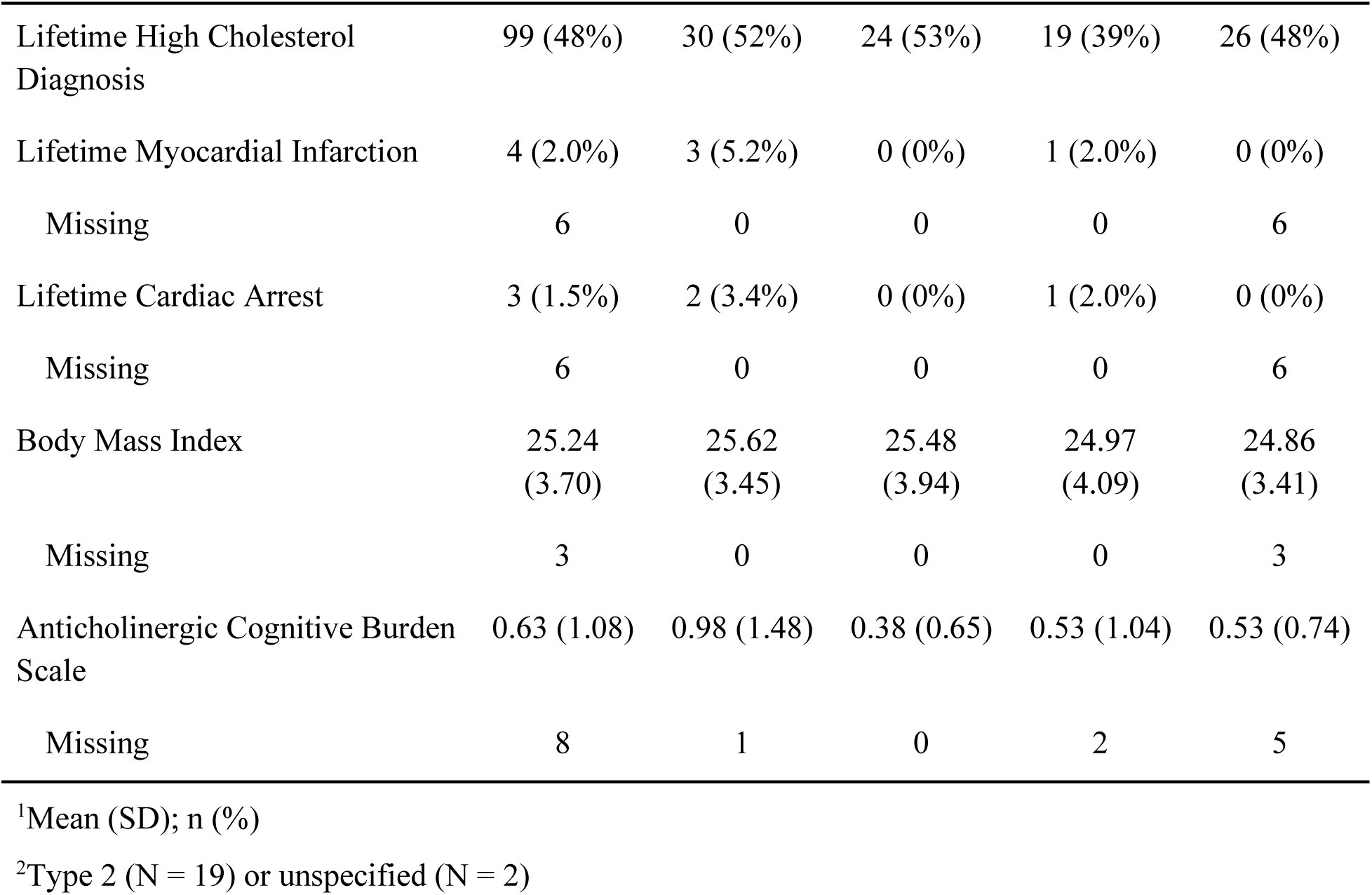
Demographic characteristics and self-reported medical history of MBAR participants, unimputed data (N = 206).

| **Characteristic** | **Overall**<br>**N = 206<sup>1</sup>** | **Site 1**<br>**N = 58<sup>1</sup>** | **Site 2**<br>**N = 45<sup>1</sup>** | **Site 3**<br>**N = 49<sup>1</sup>** | **Site 4**<br>**N = 54<sup>1</sup>** |
| --- | --- | --- | --- | --- | --- |
| Age | 88.44<br>(3.13) | 88.55<br>(3.45) | 88.49<br>(2.88) | 89.12<br>(3.43) | 87.65<br>(2.53) |
| Female | 112 (54%) | 29 (50%) | 22 (49%) | 29 (59%) | 32 (59%) |
| Race |  |  |  |  |  |
| Asian | 2 (1.0%) | 0 (0%) | 0 (0%) | 1 (2.0%) | 1 (1.9%) |
| Black/African American | 6 (2.9%) | 4 (6.9%) | 0 (0%) | 0 (0%) | 2 (3.7%) |
| Other | 1 (0.5%) | 0 (0%) | 0 (0%) | 1 (2.0%) | 0 (0%) |
| White | 197 (96%) | 54 (93%) | 45 (100%) | 47 (96%) | 51 (94%) |
| Hispanic/Latino | 6 (2.9%) | 0 (0%) | 2 (4.4%) | 0 (0%) | 4 (7.4%) |
| Years of Education | 16.11<br>(2.98) | 15.90<br>(2.67) | 16.09<br>(2.87) | 16.29<br>(3.35) | 16.19<br>(3.11) |
| MoCA Score | 24.77<br>(2.49) | 24.47<br>(2.11) | 24.36<br>(2.53) | 24.94<br>(2.82) | 25.28<br>(2.47) |
| Missing | 1 | 0 | 0 | 0 | 1 |
| Smoking Pack-Years |  |  |  |  |  |
| Nonsmoker | 94 (49%) | 26 (46%) | 18 (41%) | 21 (49%) | 29 (62%) |
| High | 47 (25%) | 15 (26%) | 13 (30%) | 11 (26%) | 8 (17%) |
| Low | 50 (26%) | 16 (28%) | 13 (30%) | 11 (26%) | 10 (21%) |
| Missing | 15 | 1 | 1 | 6 | 7 |
| Lifetime Alcohol Abuse | 2 (1.0%) | 0 (0%) | 0 (0%) | 2 (4.1%) | 0 (0%) |
| Missing | 4 | 0 | 0 | 0 | 4 |
| Lifetime Diabetes Diagnosis<sup>2</sup> | 21 (10%) | 9 (16%) | 5 (11%) | 5 (10%) | 2 (3.7%) |
| Lifetime Hypertension Diagnosis | 120 (59%) | 36 (62%) | 25 (56%) | 32 (65%) | 27 (54%) |
| Missing | 4 | 0 | 0 | 0 | 4 |
| Lifetime High Cholesterol<br>Diagnosis | 99 (48%) | 30 (52%) | 24 (53%) | 19 (39%) | 26 (48%) |
| Lifetime Myocardial Infarction | 4 (2.0%) | 3 (5.2%) | 0 (0%) | 1 (2.0%) | 0 (0%) |
| Missing | 6 | 0 | 0 | 0 | 6 |
| Lifetime Cardiac Arrest | 3 (1.5%) | 2 (3.4%) | 0 (0%) | 1 (2.0%) | 0 (0%) |
| Missing | 6 | 0 | 0 | 0 | 6 |
| Body Mass Index | 25.24<br>(3.70) | 25.62<br>(3.45) | 25.48<br>(3.94) | 24.97<br>(4.09) | 24.86<br>(3.41) |
| Missing | 3 | 0 | 0 | 0 | 3 |
| Anticholinergic Cognitive Burden<br>Scale | 0.63 (1.08) | 0.98 (1.48) | 0.38 (0.65) | 0.53 (1.04) | 0.53 (0.74) |
| Missing | 8 | 1 | 0 | 2 | 5 |

<sup>1</sup>Mean (SD); n (%)
<sup>2</sup>Type 2 (N = 19) or unspecified (N = 2)

Coefficient of variation (CoV) for a subset of the measures included in the UDS 3.0 (Table 1) was used as an index of cognitive dispersion. Greater cognitive dispersion has been argued to reflect failure of top-down executive control mechanisms (Tractenberg and Pietrzak, 2011) and may have prognostic value for future cognitive decline (Gleason et al., 2017). CoV was calculated using the demographically-corrected norms for this composite reported by Kiselica and colleagues (Kiselica et al., 2024). Norms are corrected for age, years of education, sex, and race/ethnicity (non-Hispanic White/other) and are available from 50-101 years of age (Kiselica et al., 2024).

Participants additionally completed the NIH Toolbox Cognition Battery. Crystallized-fluid cognition discrepancy was used as a proxy for cognitive decline (O’Shea et al., 2018). NIH Toolbox Fluid Cognition Composite was subtracted from NIH Toolbox Crystallized Cognition Composite to calculate crystallized-fluid discrepancy (Iverson et al., 2023), a metric of decline from estimated premorbid intellect. Because age-corrected NIH Toolbox Cognition Battery norms are not available for individuals older than 85 (Casaletto et al., 2015), uncorrected Standard Scores were converted to uncorrected z-scores (to maintain consistency with other cognitive summary scores) and used for analyses of NIH Toolbox data.

### 2.4 Structural Image Acquisition and Processing

T1-weighted MP-RAGE images were acquired on a Siemens Prisma or Skyra 3T scanner, using a 64-channel head coil, according to a standard protocol across sites (TE: 3.37ms; TR: 2530ms; flip angle = 7°; FoV = 240 x 256 x 176mm; voxel size: 1.0x1.0x1.0mm^3^; scan duration: 6 minutes and 3 seconds). Images were quality-checked visually by a trained rater. Images were then submitted to the automated brainageR 2.1 pipeline (Cole, 2019). Images were segmented into gray matter, white matter, and CSF, normalized in SPM12 (Friston et al., 2006), and visually quality-checked again; segmented GM, WM, and CSF images were then vectorized in R 4.2.2, masked to the brainageR template, and submitted to brainageR’s PCA rotation matrix. The 435 principal components produced by rotation were then entered as predictors into a Gaussian Process regression model using kernlab. Model output was brain-predicted age. BrainPAD was calculated as brain-predicted age – chronological age. No deviations were made from the default brainageR pipeline.

### 2.5 Statistical Analysis

All analyses were conducted in R 4.3.1 (R Core Team, 2022). Distribution of clinical and demographic variables of interest was reported descriptively by study site. Pearson correlation between brainPAD and chronological age was reported. To address bias created by missing data or poor-quality MR images, covariate imbalance between complete and incomplete cases was assessed descriptively using standardized mean differences (continuous variables) and raw differences in proportion (categorical variables) in the cobalt R package (Greifer, 2024); a threshold of 0.10 was used as a cutoff for imbalance. Due to the high proportion of incomplete cases (42%), incomplete cases were imputed under fully-conditional specification using mice 3.17.0 (van Buuren and Groothuis-Oudshoorn, 2011). Full details on the imputation procedure are reported in Supplementary Methods. Lifetime exercise and pack-years were categorized separately for each imputation.

To address our first aim, brainPAD was entered as the outcome of a linear model. A random site-specific intercept was initially fit in lmerTest (Kuznetsova et al., 2017) to address site effects; however, the random intercepts were dropped from analysis of imputed data due to explained variance approaching zero (i.e., no substantial variance explained by site-specific intercepts). Fixed predictor terms were age, sex, years of education, cigarette pack-year history (nonsmoker/< median/≥ median), exercise quantile, history of hypercholesterolemia diagnosis, history of hypertension diagnosis, BMI, and anticholinergic medication burden. Diabetes, myocardial infarction, cardiac arrest, Hispanic ethnicity, non-White race, and lifetime alcohol abuse were dropped as predictors due to low frequency in the sample (≤10% each in unimputed data). Parameters of interest were estimated separately for individual imputed datasets and then pooled using Rubin’s rules. Supplementary analyses were conducted using listwise deletion to assess the impact of the imputation approach on observed results; the site-specific random intercept was retained for this supplementary model, as it explained substantial variance in unimputed data.

To address our second aim, brainPAD-cognition associations were evaluated with linear mixed effects models fit in lmerTest (Kuznetsova et al., 2017). Separate models were fit with each cognitive summary score (global cognitive z-score, CoV, and fluid-crystallized discrepancy) as dependent variables. BrainPAD and age were entered as fixed predictor terms in all models. Due to the use of uncorrected scores for NIH Toolbox Fluid-Crystallized Discrepancy, sex and years of education were included as covariates in this model only. Separate random intercepts were fit for each study site. Again, parameters of interest were estimated separately for individual imputed datasets and then pooled using Rubin’s rules. Supplementary analyses were conducted using a listwise deletion approach to assess the impact of imputation on results.

## 3. RESULTS

### 3.1 Clinical and Demographic Variables

Clinical and demographic variables are reported by site in Table 1 (unimputed data). Participants were predominantly non-Hispanic White, with a mean of 16.11 (SD: 2.98) years of education. Forty-five percent of participants were male.

### 3.2 Missingness

Prior to imputation, 157 (76%) participants had interpretable brainPAD data and 119 (58%) had complete data for all variables of interest. Missingness is reported in full in Supplementary Table 2. Missingness was associated with study site, lower MoCA score, greater BMI, and greater anticholinergic burden (Supplementary Table 3).

### 3.3 BrainPAD

In unimputed data, the Pearson correlation between brain-predicted age and chronological age was 0.21 (95% CI = 0.05, 0.35, p = 0.009). The mean brainPAD was -7.99 (SD: 5.37; range: - 24.50, 6.03; Figure 1). The distribution of brainPAD by site is plotted in Figure 2.

**Figure 1.**
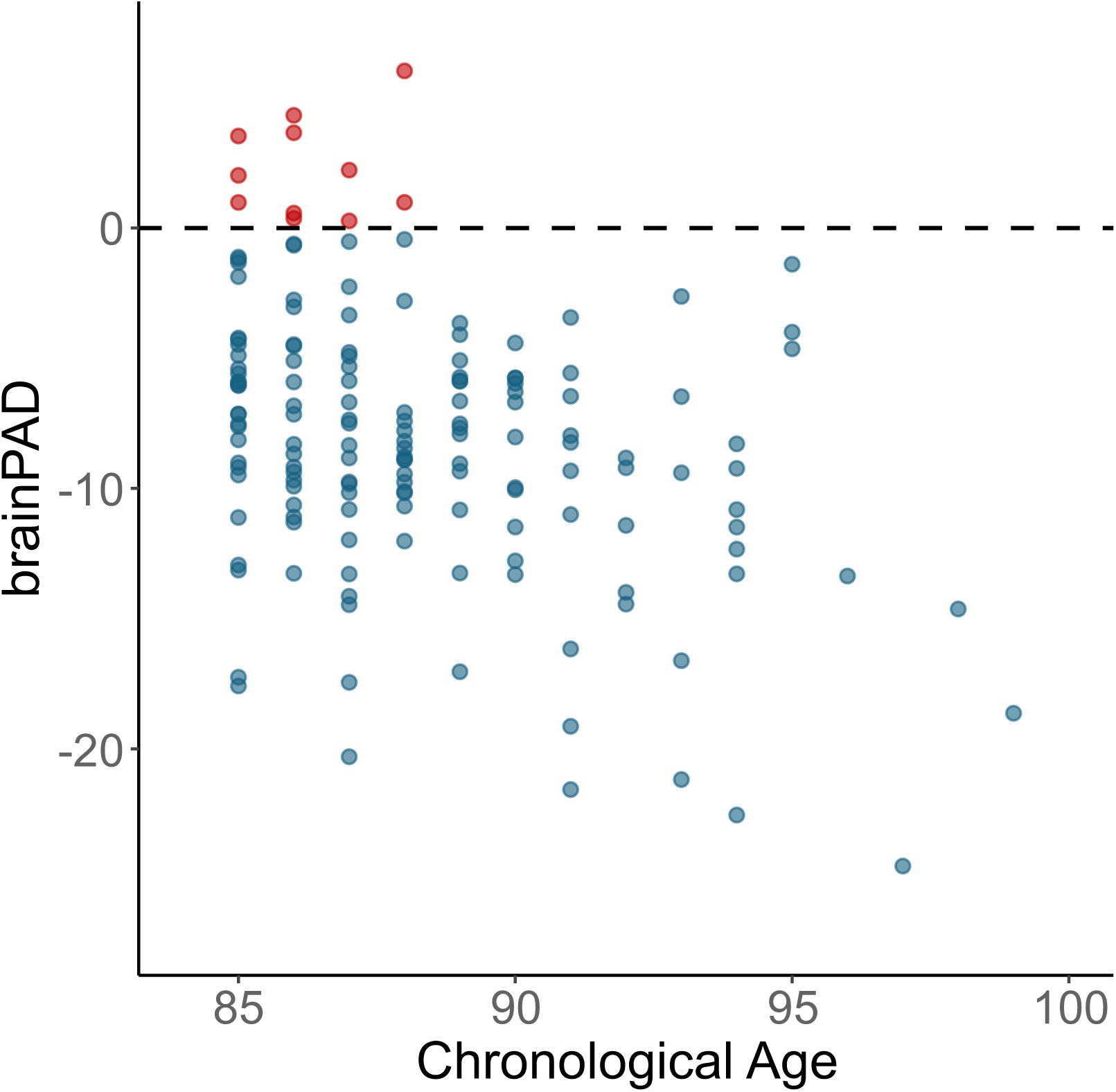
Visualization of brainPAD and chronological age in cognitively-intact oldest-old adults, unimputed data (N = 157). Mean brainPAD was -7.99 (SD: 5.37, range: -24.50, 6.03), representing a roughly 8-year delay in brain aging.

**Figure 2.**
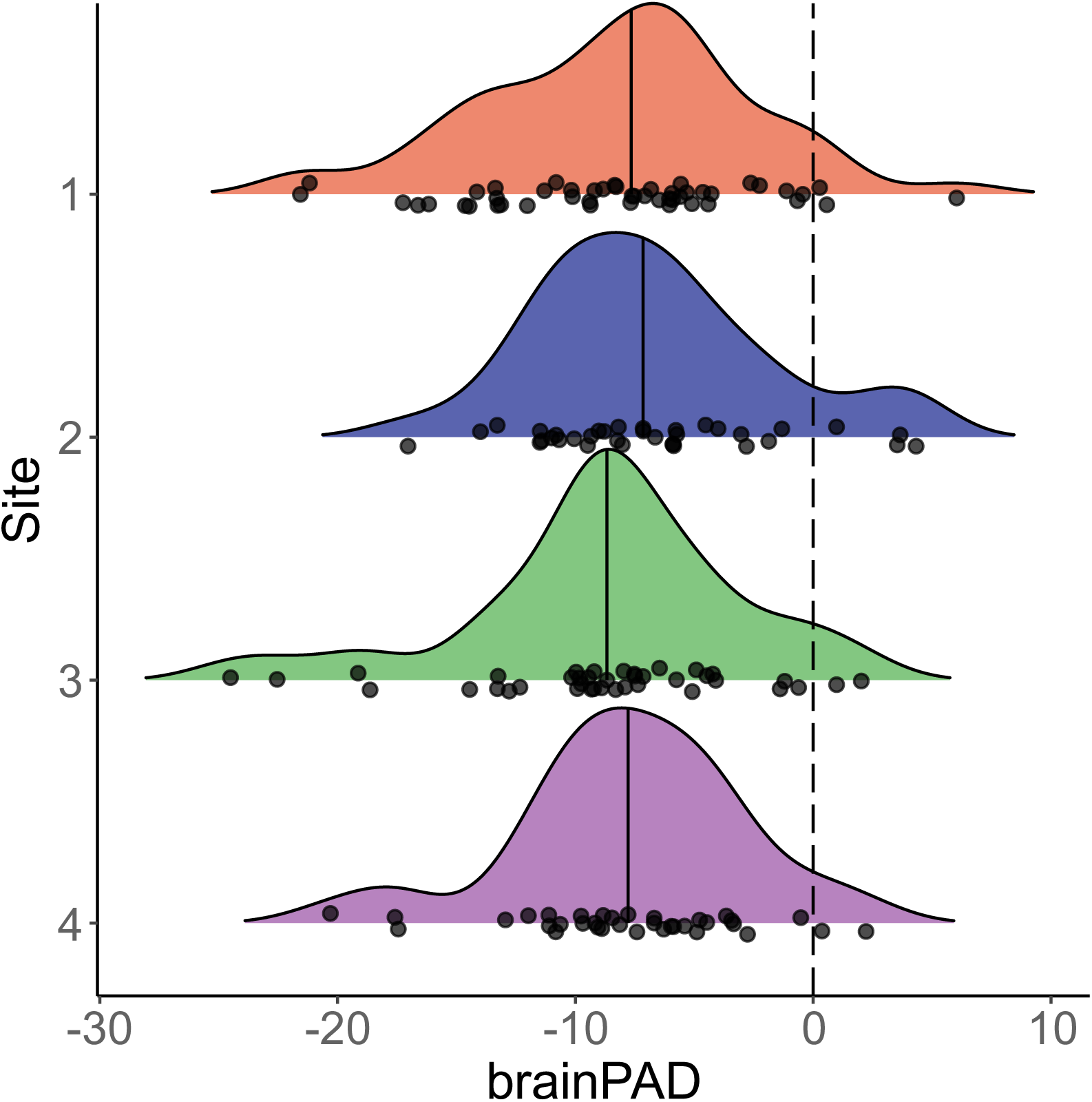
Distribution of brainPAD by study site in cognitively-intact oldest-old adults (85+), unimputed data (N = 157).

### 3.4 Lifetime Exposure History and BrainPAD

Results of linear mixed effects models on imputed data are reported in Table 3. Female sex was significantly associated with more-negative brainPAD (B = -2.35, 95% CI = -4.28, -0.41, p = 0.018; Figure 3); that is, female participants had relatively younger-appearing brains (mean: - 9.42; SD: 4.90) than males (mean: -6.34; SD: 5.43). Age was inversely associated with brainPAD (B = -0.67, 95% CI = -0.93, -0.42, p < 0.001), such that older participants showed comparatively-delayed brain aging. No other predictor variables were significantly associated with brainPAD (Table 3). Results were substantively similar in complete-case analyses (Supplementary Table 3).

**Figure 3.**
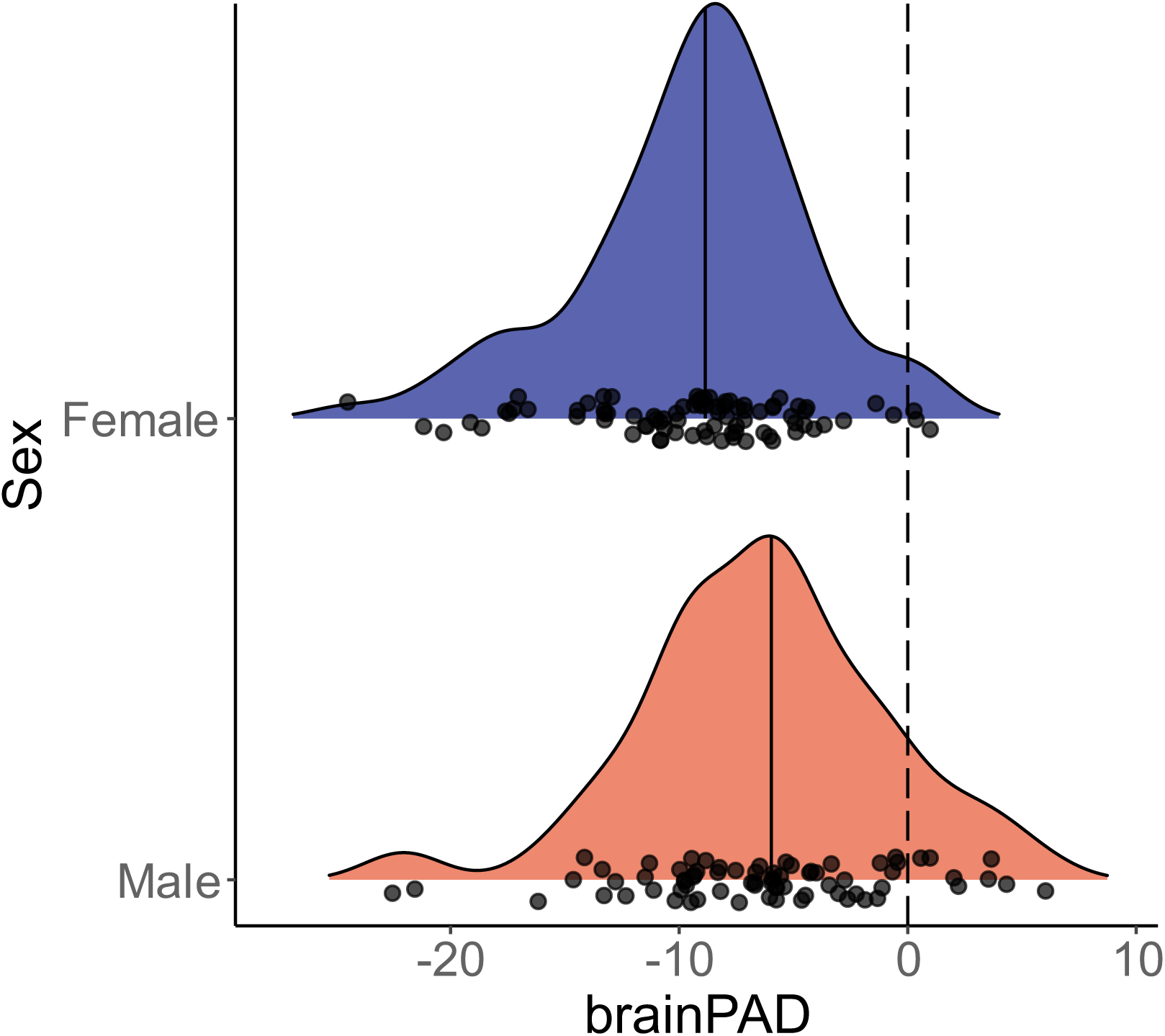
Among cognitively-intact oldest-old adults (85+), brainPAD was significantly more negative in females, unimputed data (N = 157

**Table 3.** Results of linear model predicting brainPAD from participant characteristics (N = 206), imputed data. Site-specific random intercept terms were dropped due to low variance explained.

| Characteristic | B | 95% CI<sup>1</sup> | p-value |
| --- | --- | --- | --- |
| Age | -0.67 | -0.95, -0.41 | < 0.001 |
| Female | -2.35 | -4.28, -0.41 | 0.018 |
| Smoking Pack-Years |  |  |  |
| Nonsmoker | — | — |  |
| High | -0.67 | -2.68, 1.34 | 0.51 |
| Low | -0.75 | -2.65, 1.15 | 0.43 |
| Anticholinergic Medication Use |  |  |  |
| No | — | — |  |
| Yes | -0.60 | -2.28, 1.08 | 0.48 |
| Cumulative Lifetime Exercise Quartile |  |  |  |
| Q1 | — | — |  |
| Q2 | -0.66 | -2.86, 1.53 | 0.55 |
| Q3 | -0.31 | -2.91, 2.29 | 0.81 |
| Q4 | 0.08 | -2.43, 2.60 | 0.95 |
| Lifetime Hypertension Diagnosis |  |  |  |
| No | — | — |  |
| Yes | 0.34 | -1.30, 1.98 | 0.68 |
| Lifetime Hypercholesterolemia Diagnosis |  |  |  |
| No | — | — |  |
| Yes | -0.33 | -2.08, 1.41 | 0.71 |
| Years of Education | 0.06 | -0.21, 0.33 | 0.64 |
| BMI | 0.08 | -0.18, 0.34 | 0.55 |

| Characteristic | B | 95% CI<sup>1</sup> | p-value |
| --- | --- | --- | --- |

<sup>1</sup>CI = Confidence Interval

### 3.5 BrainPAD and Cognitive Performance

The distribution of UDS 3.0 global z-score, UDS 3.0 CoV, and NIH Toolbox Crystallized-Fluid Discrepancy Score by site is visualized in Figure 4. In unimputed data, the mean UDS 3.0 global z-score was -0.26 (SD: 0.49); the mean UDS 3.0 CoV was -0.26 (SD: 1.04); and the mean NIH Toolbox Crystallized-Fluid Discrepancy z-score was 2.23 (SD: 0.69).

Results of linear mixed effects models on imputed data are reported in Table 4. There was no statistically-significant association between brainPAD and UDS 3.0 global z-score (B = -0.005, 95% CI = -0.02, 0.01, p = 0.48) or UDS 3.0 CoV (B = 0.01, 95% CI = -0.02, 0.04, p = 0.51) in models adjusted for age. There was no statistically-significant association between brainPAD and NIH Toolbox Crystallized-Fluid Discrepancy (B = 0.004, 95% CI = -0.02, 0.03, p = 0.75) in models adjusted for age, sex, and years of education. Results were substantively similar in complete-case analyses (Supplementary Table 4).

**Figure 4.**
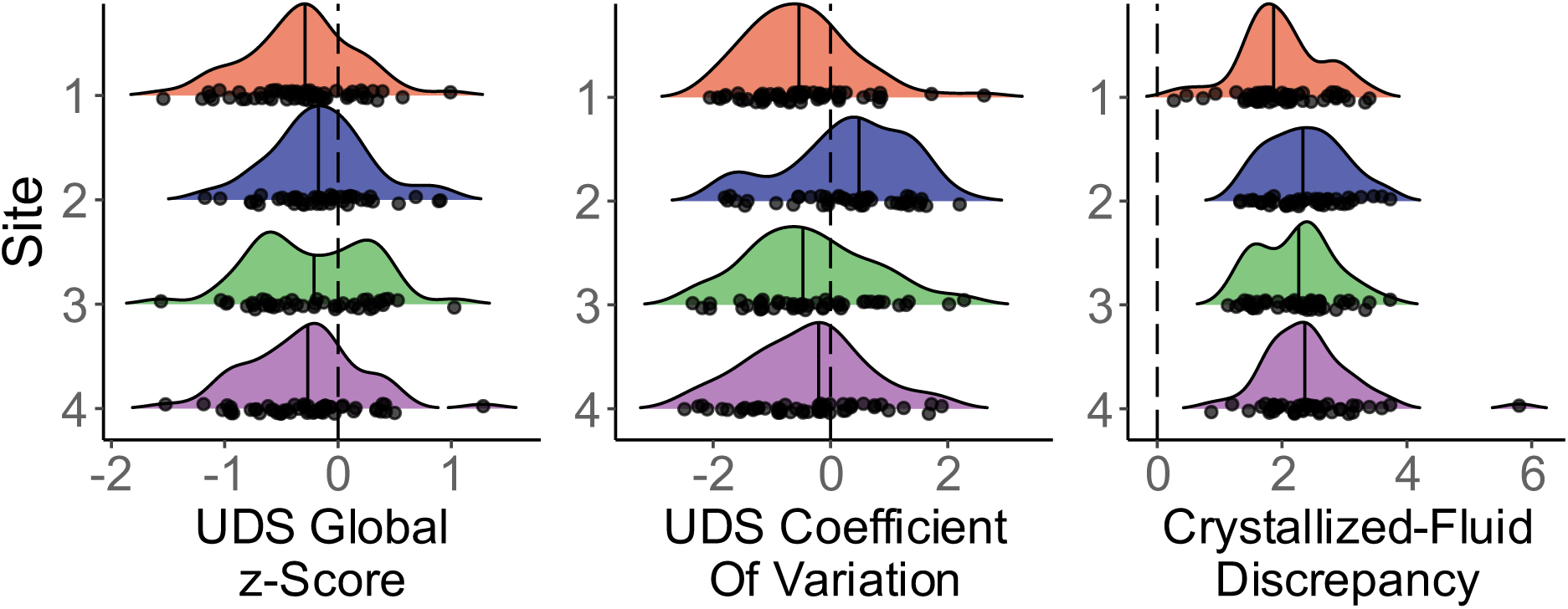
Distribution of UDS 3.0 and NIH Toolbox cognitive summary variables by study site, unimputed data. Variables are reported as z-scores.

**Table 4.** Fixed effects from linear mixed effects models of brainPAD and cognitive performance (N = 206), imputed data.

| Characteristic | UDS 3.0 Global z-score<sup>2</sup> |  |  | UDS 3.0 Coefficient of Variation<sup>3</sup> |  |  | NIH Toolbox Crystallized-Fluid Discrepancy<sup>4</sup> |  |  |
| --- | --- | --- | --- | --- | --- | --- | --- | --- | --- |
|  | B | 95% CI<sup>1</sup> | p-value | B | 95% CI<sup>1</sup> | p-value | B | 95% CI<sup>1</sup> | p-value |
| Age | -0.007 | -0.03, 0.02 | 0.58 | 0.01 | -0.04, 0.06 | 0.77 | 0.03 | -0.004, 0.65 | 0.08 |
| brainPAD | -0.005 | -0.02, 0.01 | 0.48 | 0.01 | -0.02, 0.04 | 0.51 | 0.004 | -0.02, 0.03 | 0.75 |
| Years of Education | — | — | — | — | — | — | 0.05 | 0.02, 0.09 | 0.004 |
| Female | — | — | — | — | — | — | 0.09 | -0.13, 0.31 | 0.42 |

<sup>1</sup>CI = Confidence Interval
<sup>2</sup>Adjusted for age, years of education, and sex using published norms
<sup>3</sup>Adjusted for age, years of education, sex, and race/ethnicity using published norms
<sup>4</sup>Uncorrected scores were used due to unavailability of age-corrected NIH Toolbox norms for individuals over 85

## 4. DISCUSSION

Although brainPAD is a promising biomarker of age-related differences in brain structure, it has not been extensively examined in oldest-old adults to date. Cross-sectional associations between exposure history, biomarkers of aging, and cognitive performance may not generalize from young-old to oldest-old samples due to nonlinear age effects and survivorship effects; survivorship effects may be particularly pronounced in cognitively-intact oldest-old individuals. We have previously referred to this effect as “surviving and thriving” (Britton et al., 2025). The present analysis examined associations between self-reported exposure history and brainPAD, as well as brainPAD and three cognitive summary scores, in a cross-sectional sample of cognitively-intact oldest-old adults.

Overall, participants showed structural brain features consistent with delayed brain aging: the mean brainPAD in unimputed data was -7.99 (SD: 5.37) across all participants, and the mean in females was -9.42 (SD: 4.90). This finding parallels results recently reported by Park and colleagues (Park et al., 2025), who observed delayed brain aging in a sample of young-old and oldest-old “superagers”. Collectively these results suggest that some cognitively-intact aging adults may be physiologically younger than birth cohort peers, consistent with a “brain maintenance” model of healthy cognitive aging (Cabeza et al., 2018; Merenstein and Bennett, 2022).

While we are unable to conclusively identify specific protective factors in our cross-sectional observational sample, delayed brain aging may be related to protective health behavior or cognitive reserve. Our participants were atypically healthy members of their birth cohort. The lifetime prevalence of myocardial infarction, cardiac arrest, diabetes mellitus, alcohol abuse or dependence, and smoking in our sample was low compared to representative population-based US estimates from comparable birth cohorts (Anderson et al., 2012; Bishop et al., 2022; Fishman et al., 2014; Grant, 1997). The mean educational attainment in the sample was 16.29 years (SD: 3.06), consistent with high adult SES and high cognitive reserve (although it should be noted that associations between educational attainment and brain-predicted age are inconsistent) (Nyberg et al., 2021; Steffener et al., 2016; Wagen et al., 2022). In sum, our sample of healthy and cognitively-intact oldest-old participants may represent a best-case scenario for brain function in aging.

Within this highly-selected sample of cognitively-intact oldest-old adults, brainPAD was not associated with lifetime exposure history. Self-reported pack-year history, hypertension, hypercholesterolemia, cumulative lifetime exercise, BMI, and anticholinergic medication burden were not significantly associated with brainPAD. These findings contrast with prior reports of statistically-significant associations between smoking (Bittner et al., 2021; Linli et al., 2022), cardiovascular disease (Cherbuin et al., 2021; De Lange et al., 2020; Wagen et al., 2022), and physical fitness (Bittner et al., 2021; Dunås et al., 2021; Steffener et al., 2016) and greater brainPAD in midlife and young-old samples, as well as reported associations between medication burden and aging more generally. However, our findings parallel growing evidence that risk factors such as diabetes, APOE4, hypertension, and obesity may have attenuated apparent effects on cognitive status and mortality in oldest-old samples (Bemmel et al., 2006; Corrada et al., 2017, 2013, 2008; Juva et al., 2000; Luchsinger et al., 2007), or in some samples may even appear protective (Corrada et al., 2017; Oates et al., 2007).

Attenuation of exposure-outcome associations in oldest-old adults has been attributed to survivorship effects. As a birth cohort ages, the least-healthy individuals are disproportionately lost from the cognitively-intact population due to earlier death or incident cognitive impairment. Consequently, surviving oldest-old adults are heavily selected both for overall health and for low susceptibility to common risk factors (e.g., smoking or hypertension): for instance, surviving smokers may be particularly resistant to the adverse physiological impact of smoking. Therefore, exposures harmful on the population level may appear neutral or even beneficial in the oldest-old due to disproportionate attrition of susceptible individuals prior to sampling (Glymour, 2007; Hernán et al., 2008; Liu et al., 2010). This effect, although apparent even in population-based samples of the oldest-old, may be particularly pronounced in cognitively-intact oldest-old adults with high-quality MR data (i.e., the individuals represented in imaging studies of healthy aging). While our findings are generally consistent with survivorship effects, it should also be noted that brainPAD derived from T1-weighted MR images may not fully capture the effects of cardiovascular risk factors on white matter hyperintensity volumes, subclinical infarcts, and microbleeds; replication using multimodal brainPAD would be informative (Cole, 2020).

Notably, female sex was significantly associated with more-negative brainPAD (i.e., a younger-looking brain). Sex differences in brainPAD have previously been reported; however, the direction of these effects is inconsistent, with some studies identifying more-positive brainPAD in females (Boyle et al., 2021; Smith et al., 2019), others identifying more-positive brainPAD or faster longitudinal increase in brainPAD in males (Cole et al., 2018; Franke et al., 2013; Wagen et al., 2022; Wrigglesworth et al., 2022a), and others finding no sex difference (Bittner et al., 2021). Discrepant results may be partially explained by distinct risk factors for brain aging by sex. Franke and coauthors report that brain-predicted age is associated with metabolic markers in aging males, but not females (Franke et al., 2014); in young adults, Sanford and coauthors likewise report sex-related differences in variables associated with brain-predicted age (Sanford et al., 2022). However, in oldest-old adults, sex-related differences may also be explained by discrepant survivorship and the “health-survival paradox.” Survivorship effects act differentially on men and women: in high-income societies, women typically have longer life expectancies but shorter disability-free life expectancies (Hoogendijk et al., 2019; Oksuzyan et al., 2008). Indeed, we have previously reported sex-dependent age-GABA associations in this sample, potentially due to more stringent survivorship effects in oldest-old males (Britton et al., 2025). Overall, our findings underline that sex is likely relevant to brainPAD and, in combination with prior work in this area, suggest that sex effects may be birth cohort-specific.

The second aim of our analysis was to evaluate associations between brainPAD and three summary measures of cognitive function in oldest-old adults. Prior studies have reported associations between more-positive brainPAD and worse cognition in lifespan samples (Boyle et al., 2021; Liem et al., 2017; Yin et al., 2023), midlife samples (DeJong et al., 2024; Elliott et al., 2021), and typical aging samples (Cole et al., 2018; Jawinski et al., 2022; Park et al., 2025; Wagen et al., 2022; Wrigglesworth et al., 2022b). While brainPAD-cognition associations have not always been replicated (Boyle et al., 2021; Tetereva and Pat, 2024) and causality may be temporally complex (i.e., poor early-life cognition has been associated with more-positive brainPAD in adulthood) (Elliott et al., 2021), brainPAD appears to have predictive value for future cognitive decline in at-risk aging adults (Biondo et al., 2022; Gaser et al., 2013). Consequently, brainPAD has been interpreted as a potentially clinically-meaningful biomarker of cognitive aging (Cole et al., 2019).

In our sample, brainPAD was not associated with global cognitive function, coefficient of variation, or crystallized-fluid cognitive discrepancy. This finding may be explained by several mutually-compatible mechanisms. First, the range of cognitive function within our sample was constrained by study inclusion criteria, which may have limited our ability to detect subtle associations across the range of cognitive function in aging. Second, subtle domain-specific associations may not be apparent in global summary measures of cognition, although CoV is typically sensitive to subtle cognitive changes in normal aging and neurological disease (Aita et al., 2024b; Del Bene et al., 2025; Jackson et al., 2012; Webber et al., 2024). Third, although the mean brainPAD corresponded to a roughly eight-year delay in brain aging in the sample overall, brainPAD in unimputed data ranged from -24.50 to 6.03. This broad range suggests that our sample may have included both participants resistant to age-related brain changes (i.e., effective brain maintenance) and participants with strong compensatory abilities in the presence of age-related brain changes (i.e., strong cognitive reserve) (Stern, 2009); that is, several distinct brain aging phenotypes may be represented among cognitively-intact oldest-old adults (Eavani et al., 2018), reflecting potentially dissociable profiles of risk and protective factors (Wrigglesworth et al., 2023). The association between frank neuropathological changes and cognitive function appears to be attenuated in the oldest-old (Balasubramanian et al., 2012; Middleton et al., 2011; Savva et al., 2009), consistent with strong compensatory ability in a substantial subset of non-demented oldest-old adults. Our findings, although not definitive, suggest that cognitively-intact oldest-old adults may be a physiologically-heterogeneous group whose cognitive performance may be driven by varying combinations of brain maintenance and cognitive reserve. Follow-up work in larger samples stratified by brain age or educational attainment would contribute to disentangling maintenance vs. reserve pathways in the cognitively-intact oldest-old.

Our cross-sectional results emphasize the need for prospective cohort studies of brain aging in two major ways. First, attrition due to nonrandom mortality and incident cognitive impairment can be directly observed in prospective cohort studies and the resulting biases addressed analytically (e.g., with inverse probability weighting) (Handels et al., 2020). Second, it is currently unclear to what extent brainPAD validly predicts future cognitive performance in oldest-old adults. The absence of cross-sectional brainPAD-cognition associations in our sample suggests that biomarkers of physiological aging may be comparatively less associated with cognition in oldest-old age (vs. young-old age) or that heterogeneity in this population may be greater. Balasubramanian and colleagues have previously reported that frank neuropathology is not associated with nonlinear cognitive trajectory over time in oldest-old adults, potentially due to selection for strong compensatory abilities (Balasubramanian et al., 2012). Our findings raise the possibility that brainPAD may have limited predictive power in this population; longitudinal studies of oldest-old adults with baseline brainPAD observation will be necessary to evaluate its clinical relevance.

Several limitations should be kept in mind when interpreting our findings. First, lifetime exposures were self-reported retroactively. Consequently, participants may have been misclassified due to inaccurate recall or due to social desirability bias. Additionally, historical information on BMI and medication exposure in midlife and young-old age was not available; consequently, reverse causality (e.g., weight loss due to declining health) may have masked underlying associations between exposures in midlife and brainPAD. Replication with exposure measurements across the lifespan (e.g., from health record data) would be beneficial in characterizing associations between health history and brainPAD. Furthermore, although the brain-predicted age value generated by brainageR has demonstrated predictive value in other samples (Biondo et al., 2022), multimodal imaging models may better capture some aspects of physiological aging (Cole, 2020). Finally, our sample was largely homogeneous in racial/ethnic composition and high educational attainment; replication in diverse samples would improve understanding of aging-related selection effects across subgroups (Gilsanz et al., 2019).

In conclusion, we observed a roughly 8-year delay in brain aging in a sample of cognitively-intact oldest-old adults. Within our sample, brainPAD was not associated with risk factors with well-established relationships to physiological aging at a population level, consistent with atypical resilience among individuals who have survived to oldest-old age. Our findings highlight the potential impact of population- and study-level selection effects on observed associations in aging research. From a translational perspective, our study emphasizes that more research is needed to identify factors that contribute to the survival and resilience of cognitively-intact oldest-old individuals, as these factors may shed light on personalized approaches to the deceleration of brain aging.

## Author Contributions

**Conceptualization:** Mark K. Britton, Clinton B. Wright, David A. Raichlen, Victor A. Del Bene, Virginia G. Wadley, Theodore P. Trouard, Noam Alperin, Bonnie E. Levin, Tatjana Rundek, Kristina M. Visscher, Gene E. Alexander, Ronald A. Cohen, Eric C. Porges, and Joseph M. Gullett.

**Data curation:** Mark K. Britton, Hannah Hoogerwoerd, Joshua Juhasz, Keyanni J. Johnson, Paul D. Stewart, and Stacy S. Merritt.

**Formal analysis:** Mark K. Britton.

**Funding acquisition:** Clinton B. Wright, David A. Raichlen, Virginia G. Wadley, Theodore P. Trouard, Noam Alperin, Bonnie E. Levin, Tatjana Rundek, Kristina M. Visscher, Gene E. Alexander, and Ronald A. Cohen.

**Investigation:** Mark K. Britton, Hannah Hoogerwoerd, Joshua Juhasz, Keyanni J. Johnson, Paul D. Stewart, Stacy S. Merritt, Cortney J. Jessup, Clinton B. Wright, David A. Raichlen, G. Alex Hishaw, Virginia G. Wadley, Theodore P. Trouard, Noam Alperin, Bonnie E. Levin, Tatjana Rundek, Kristina M. Visscher, Gene E. Alexander, and Ronald A. Cohen.

**Methodology:** Mark K. Britton, Hannah Hoogerwoerd, Clinton B. Wright, David A. Raichlen, G. Alex Hishaw, Victor A. Del Bene, Virginia G. Wadley, Theodore P. Trouard, Noam Alperin, Bonnie E. Levin, Tatjana Rundek, Kristina M. Visscher, Gene E. Alexander, Ronald A. Cohen, Eric C. Porges, and Joseph M. Gullett.

**Project administration:** Paul D. Stewart, Stacy S. Merritt, and Cortney J. Jessup.

**Software:** Mark K. Britton.

**Supervision:** Eric C. Porges and Joseph M. Gullett.

**Visualization:** Mark K. Britton.

**Writing - original draft:** Mark K. Britton, Hannah Hoogerwoerd, and Joshua Juhasz.

**Writing - review & editing:** Clinton B. Wright, David A. Raichlen, G. Alex Hishaw, Victor A. Del Bene, Virginia G. Wadley, Theodore P. Trouard, Noam Alperin, Bonnie E. Levin, Tatjana Rundek, Kristina M. Visscher, Gene E. Alexander, Ronald A. Cohen, Eric C. Porges, and Joseph M. Gullett.

## Data Statement

All code used in this analysis is available from the Open Science Framework at https://osf.io/ax93d. Data are available from the McKnight Brain Research Foundation upon reasonable request.

