## Supplementary Materials for "Your Brain Doesn’t Look a Day Past 70! Cross-Sectional Associations with Brain-Predicted Age in the Cognitively-Intact Oldest-Old"

Supplementary Table 1. Covariate imbalance between complete and incomplete cases. Standardized mean differences (continuous) or raw differences in proportion (categorical) are bolded at an absolute threshold of ≥0.10.

| **Characteristic** | **Complete**  N = 119^1^ | **Incomplete**  N = 87^1^ | **SMD**^2^ |
| --- | --- | --- | --- |
| Site |  |  |  |
| 1 | 37 (31%) | 21 (24%) | -0.07 |
| 2 | 33 (28%) | 12 (14%) | **0.14** |
| 3 | 25 (21%) | 24 (28%) | 0.07 |
| 4 | 24 (20%) | 30 (34%) | **0.14** |
| Age | 88.30 (3.20) | 88.62 (3.04) | **0.10** |
| Female | 64 (54%) | 48 (55%) | -0.01 |
| Race |  |  |  |
| Asian | 0 (0%) | 2 (2.3%) | 0.02 |
| Black/African American | 6 (5.0%) | 0 (0%) | -0.05 |
| Other | 1 (0.8%) | 0 (0%) | -0.01 |
| White | 112 (94%) | 85 (98%) | 0.04 |
| Hispanic/Latino | 4 (3.4%) | 2 (2.3%) | 0.01 |
| Years of Education | 16.34 (3.00) | 15.79 (2.95) | **-0.18** |
| MoCA Score | 24.91 (2.65) | 24.57 (2.25) | **-0.14** |
| Lifetime Alcohol Abuse | 1 (0.8%) | 1 (1.2%) | 0.004 |
| Smoking Pack-Years |  |  |  |
| Nonsmoker | 61 (51%) | 33 (46%) | -0.05 |
| High | 25 (21%) | 22 (31%) | **0.10** |
| Low | 33 (28%) | 17 (24%) | -0.04 |
| Lifetime Hypertension Diagnosis | 70 (59%) | 50 (60%) | 0.01 |
| Lifetime High Cholesterol Diagnosis | 59 (50%) | 40 (46%) | -0.04 |
| Lifetime Diabetes Diagnosis | 14 (12%) | 7 (8.0%) | -0.04 |
| Lifetime Myocardial Infarction | 3 (2.5%) | 1 (1.2%) | -0.01 |
| Lifetime Cardiac Arrest | 2 (1.7%) | 1 (1.2%) | -0.005 |
| Body Mass Index | 24.84 (3.21) | 25.82 (4.26) | **0.26** |
| Cumulative Lifetime Exercise Quartile |  |  |  |
| Q1 | 33 (28%) | 7 (17%) | **-0.11** |
| Q2 | 29 (24%) | 11 (27%) | 0.02 |
| Q3 | 32 (27%) | 8 (20%) | -0.07 |
| Q4 | 25 (21%) | 15 (37%) | 0.16 |
| Anticholinergic Cognitive Burden Scale | 0.57 (1.03) | 0.71 (1.15) | **0.13** |
| Brain-Predicted Age | 80.44 (5.43) | 80.07 (3.80) | -0.08 |
| ^1^n (%); Mean (SD) | | | |
| ^2^Standardized mean difference (continuous); raw difference in proportions (categorical) | | | |

Supplementary Table 2. Percentage of complete cases and reason for missingness (where applicable) for variables of interest.

| **Characteristic** | **N (%)** | **Reason for Missingness** |
| --- | --- | --- |
| Age | 206 (100%) | — |
| Sex | 206 (100%) | — |
| Race | 206 (100%) | — |
| Hispanic/Latino | 206 (100%) | — |
| Years of Education | 206 (100%) | — |
| MoCA Score | 205 (99.5%) | Incomplete MoCA data (N = 1). |
| Smoking Pack-Years | 191 (93%) | One or both of smoking duration (N = 11) or # of cigarettes per day (N = 11) not reported |
| Lifetime Alcohol Abuse | 202 (98%) | Not self-reported (N = 4) |
| Lifetime Diabetes Diagnosis | 206 (100%) | — |
| Lifetime Hypertension Diagnosis | 202 (98%) | Not self-reported (N = 4) |
| Lifetime High Cholesterol Diagnosis | 206 (100%) | — |
| Lifetime Myocardial Infarction | 200 (97%) | Not self-reported (N = 6) |
| Lifetime Cardiac Arrest | 200 (97%) | Not self-reported (N = 6) |
| Body Mass Index | 203 (99%) | Height and weight not collected (N = 3) |
| Exercise Quartile | 160 (78%) | Incomplete data for Lifetime Physical Activity Questionnaire (N = 45); Lifetime Physical Activity Questionnaire not returned (N = 13) |
| Anticholinergic Cognitive Burden Scale | 198 (96%) | Medications not self-reported (N = 8) |
| Brain-Predicted Age | 157 (76%) | MRI not completed (N = 31); structural MRI failed visual quality control (N = 18) |
| UDS 3.0 Global z-Score | 202 (98%) | Either or both of Trails B (N = 4) or Craft Story 21 Immediate Recall, paraphrase (N = 1) not completed |
| UDS 3.0 Coefficient of Variation | 202 (98%) | Trails B not completed (N = 4) |
| NIH Toolbox Crystallized-Fluid Discrepancy | 191 (92.7%) | One or more of NIH Toolbox Picture Sequencing (N = 14), Flanker (N = 12), Pattern Comparison (N = 12), Card Sort (N = 11), List Sorting (N = 11), Oral Reading (N = 12), or Picture Vocabulary (N = 12) not completed |

Supplementary Table 3. Fixed effects from linear mixed effects model predicting brainPAD from participant characteristics on unimputed data using a listwise deletion approach (N = 125).

| **Characteristic** | **Beta** | **95% CI**^1^ | **p-value** |
| --- | --- | --- | --- |
| Age | -0.65 | -0.95, -0.35 | <0.001 |
| Sex |  |  |  |
| Male | — | — |  |
| Female | -3.0 | -5.4, -0.69 | 0.012 |
| Smoking Pack-Years |  |  |  |
| Nonsmoker | — | — |  |
| High | -0.98 | -3.6, 1.6 | 0.5 |
| Low | -1.3 | -3.6, 0.96 | 0.2 |
| Anticholinergic Medication Use | -0.56 | -2.7, 1.5 | 0.6 |
| Cumulative Lifetime Exercise Quartile |  |  |  |
| Q1 | — | — |  |
| Q2 | -0.77 | -3.5, 2.0 | 0.6 |
| Q3 | -1.4 | -4.2, 1.5 | 0.3 |
| Q4 | -0.17 | -3.3, 3.0 | >0.9 |
| Lifetime Hypertension Diagnosis | 0.28 | -1.8, 2.4 | 0.8 |
| Lifetime Hypercholesterolemia Diagnosis | -0.96 | -3.1, 1.2 | 0.4 |
| Years of Education | 0.14 | -0.19, 0.47 | 0.4 |
| BMI | 0.12 | -0.21, 0.45 | 0.5 |
| ^1^CI = Confidence Interval | | | |

Supplementary Table 4. Fixed effects from linear mixed effects models of brainPAD and cognitive performance, unimputed data.

|  | **UDS 3.0 Global z-score^2^ (N = 156)** | | | **UDS 3.0 Coefficient of Variation^3^ (N = 156)** | | | **NIH Toolbox Crystallized-Fluid Discrepancy^4^ (N = 152)** | | |
| --- | --- | --- | --- | --- | --- | --- | --- | --- | --- |
| **Characteristic** | **B** | **95% CI**^1^ | **p-value** | **B** | **95% CI**^1^ | **p-value** | **B** | **95% CI**^1^ | **p-value** |
| Age | -0.01 | -0.04, 0.01 | 0.4 | 0.00 | -0.06, 0.06 | >0.9 | 0.04 | 0.00, 0.08 | 0.041 |
| brainPAD | 0.00 | -0.02, 0.01 | 0.7 | 0.02 | -0.01, 0.05 | 0.3 | 0.00 | -0.02, 0.02 | 0.8 |
| Years of Education | — | — | — | — | — | — | 0.05 | 0.01, 0.08 | 0.012 |
| Female | — | — | — | — | — | — | 0.03 | -0.20, 0.25 | 0.8 |
| ^1^CI = Confidence Interval ^2^Adjusted for age, years of education, and sex using published norms ^3^Adjusted for age, years of education, sex, and race/ethnicity using published norms ^4^Uncorrected scores were used due to unavailability of age-corrected NIH Toolbox norms for individuals over 85 | | | | | | | | | |

Supplementary Methods.

Multiple imputation of incomplete cases was performed using the MICE algorithm, implemented in mice 3.17.0 (van Buuren and Groothuis-Oudshoorn, 2011) and R 4.3.1 (R Core Team, 2022).

The predictor matrix was developed using the *quickpred* function, which selects predictors for each incomplete variable using Spearman correlations; we selected minpuc = 20% usable cases and mincor = 0.15 minimum rho to achieve a target of 15-25 mean predictors per incomplete variable (van Buuren, 2018). The predictor matrix included site, age, gender, and brain-predicted age as predictors for all variables; age and gender were selected due to association with brainPAD in complete-case analyses. After manual editing of the prediction matrix to break feedback loops (e.g., to prevent imputation of binarized lifetime smoking history from reported smoking duration), the mean predictors per incomplete variable was 23.94.

We ran m = 20 imputations and maxit = 20 iterations, using the default burn-in of 5000. The random forest (rf) algorithm was used for all variables. Convergence was determined by visual inspection of model trace plots.
